# Standardizing Human Brain Parcellations

**DOI:** 10.1101/845065

**Authors:** Ross M. Lawrence, Eric W. Bridgeford, Patrick E. Myers, Ganesh C. Arvapalli, Sandhya C. Ramachandran, Derek A. Pisner, Paige F. Frank, Allison D. Lemmer, Aki Nikolaidis, Joshua T. Vogelstein

**Affiliations:** Johns Hopkins University; University of Texas at Austin; Child Mind Institute

## Abstract

Using brain atlases to localize regions of interest is a requirement for making neuroscientifically valid statistical inferences. These atlases, represented in volumetric or surface coordinate spaces, can describe brain topology from a variety of perspectives. Although many human brain atlases have circulated the field over the past fifty years, limited effort has been devoted to their standardization. Standardization can facilitate consistency and transparency with respect to orientation, resolution, labeling scheme, file storage format, and coordinate space designation. Our group has worked to consolidate an extensive selection of popular human brain atlases into a single, curated, open-source library, where they are stored following a standardized protocol with accompanying metadata, which can serve as the basis for future atlases. The repository containing the atlases, the specification, as well as relevant transformation functions is available at https://github.com/neurodata/neuroparc.

## 1 Introduction

Understanding the brain’s organization is one of the key challenges in human neuroscience [1] and is critical for clinical translation [2]. Parcellation of the brain into functionally and structurally distinct regions has seen impressive advances in recent years [3], and has grown the field of network neuroscience [4, 5]. Through a range of techniques such as clustering [6–9], multivariate decomposition [10, 11], gradient based connectivity [1, 12–16], and multimodal neuroimaging [1], parcellations have enabled fundamental insights into the brain’s topological organization and network properties [17]. In turn, these properties have allowed researchers to investigate brain-behavioral associations with developmental [18, 19], cognitive [20, 21], and clinical phenotypes [22–24].

More recently, researchers interested in understanding brain organization are presented with a variety of brain atlases that can be used to define nodes of network-based analyses [25]. While this variety is a boon to researchers, the use of different parcellations across studies makes assessing reproducibility of brain-behavior relationships difficult (e.g. comparing across parcellations with different organizations and numbers of nodes; [5]). Amalgamating multiple brain parcellations into a single, standardized, curated list would offer researchers a valuable resource for evaluating replication of neuroimaging studies.

Some efforts to consolidate these atlases is already underway. For example, Nilearn is a popular Python package that provides machine-learning and informatics tools for neuroimaging [26]. Nilearn provides several single line command line interface functions to ‘fetch’ both atlases and datasets. Nilearn includes twelve anatomically and functionally defined atlases, such as the Harvard-Oxford [27] and Automated Anatomical Labeling (AAL) [28] parcellation. Although a promising prototype, Nilearn’s current atlas collection represent a limited range of available atlases, and the more recent gradient based, surface based, and multimodal parcellations have yet to be included into any central repository. More importantly, existing atlas repositories have not attempted to systematically standardize their collections following a single specification. Without well-established standards, the investigator is faced with limited information about each atlas, so connecting neuroscientific findings to the organization of the atlas becomes more difficult. Moreover, if the investigator requires a comparison across atlases, some form of metadata must be available that describes the similarities and differences between them. Neuroparc mitigates these issues by providing: (1) a detailed atlas specification which will enable researchers to both easily understand existing atlases and generate new atlases compliant with this specification, (2) a repository of the most commonly used atlases in neuroimaging, all stored in that specification, and (3) a set of functions for transforming, comparing, and visualizing these atlases. The Neuroparc package presented here includes 46 different adult human brain parcellations—including surface-based and volume-based. Here, we provide an overview of the relationship between these parcellations via comparison of the spatial similarity between atlases, as measured by Dice coefficient and adjusted mutual information. To facilitate replication and extension of this work, all the data and code are available from https://github.com/neurodata/neuroparc.

## 2 Results

### Atlases

Through the use of the python scripts provided in Neuroparc, 46 atlases were resampled to either 1 mm^3^, 2 mm^3^, or 4 mm^3^ and registered to the Montreal Neurological Institute 152 Nonlinear 6th generation reference brain (MNI). Each atlas had a accompanying JSON file containing the relevant metadata described in Methods. Of the 46 atlases, 17 lost at least one ROI from the resampling and registration process, which is recorded in the JSON files. This phenomenon was more common with atlases containing a greater amount of ROIs due to the smaller average ROI size. The smaller the ROI, the greater the chance it is overwritten by surrounding larger ROIs when down-sampled or registered.

### Dice Coefficient

A python script, provided in using functions from both AFNI [37] and FSL [38], was used to calculate the Dice Coefficients between pairs of atlases. See the Dice Coefficient section in Methods for more information about the calculation. Dice Score Maps, such as Figure 2, for each of the pairs of atlases can be found on the Neuroparc OSF repository https://osf.io/67a3t/. The purpose of these maps are to both reinforce the differences between each parcellation and what they represent, as well as serve as comparative metric for ROIs across atlases. Access to these values allow for easy cross-parcellation analysis, a useful tool to anyone using Neuroparc. On inspection, each Dice Coefficient Map was accurate in its representation of overlap present between ROIs of two different atlases.

**Table 1:**
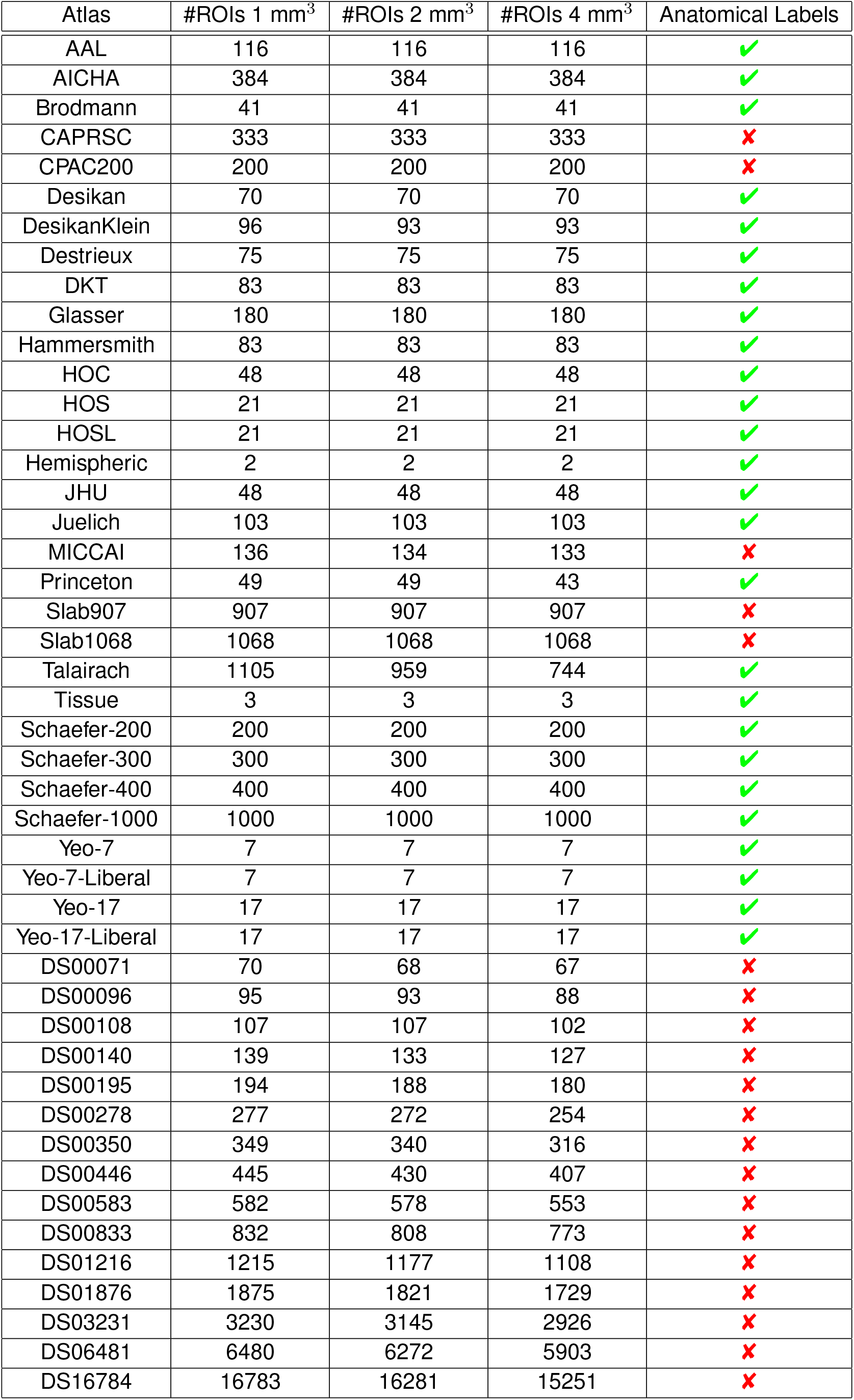
This table contains the atlases included in Neuroparc and the number of ROIs per voxel size, showing the number of ROIs lost during resampling and registration. Which atlases have anatomical labeling metadata is also noted

**Figure 1:**
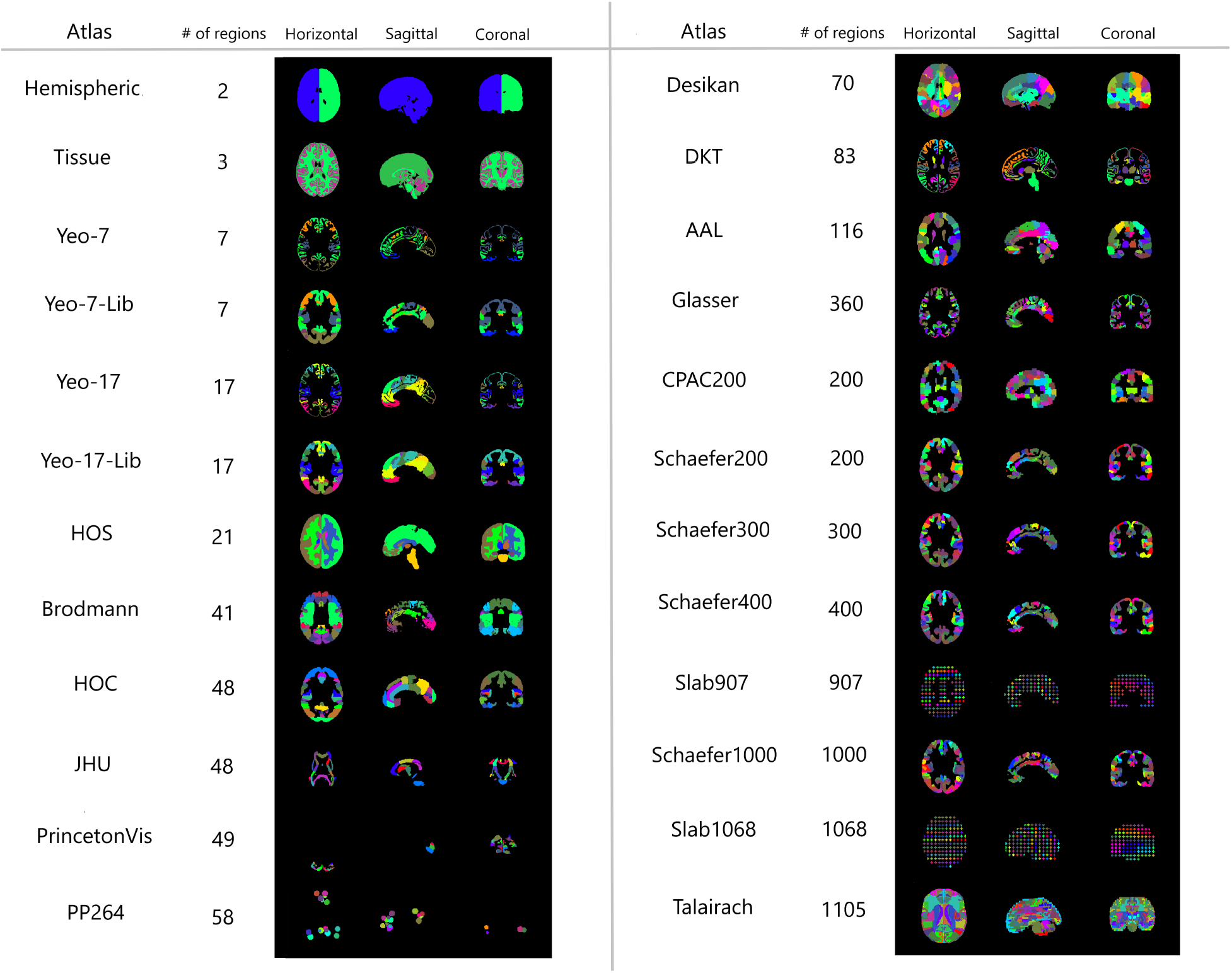
A comparison of the regions present in the major atlases available in Neuroparc. These visualizations were made using MIPAV tri-planar views on the same slice numbers. Each atlas shows a cross-section in each of the canonical orthogonal planes (H=Horizontal, S=Sagittal, C=Coronal). For most atlases, the slice numbers were (90, 108, 90). There are a few exceptions for visualization purposes: JHU: (90, 108, 109), Slab907: (95, 104, 95), Slab1068: (93, 105, 93) [1, 27–36]

**Figure 2:**
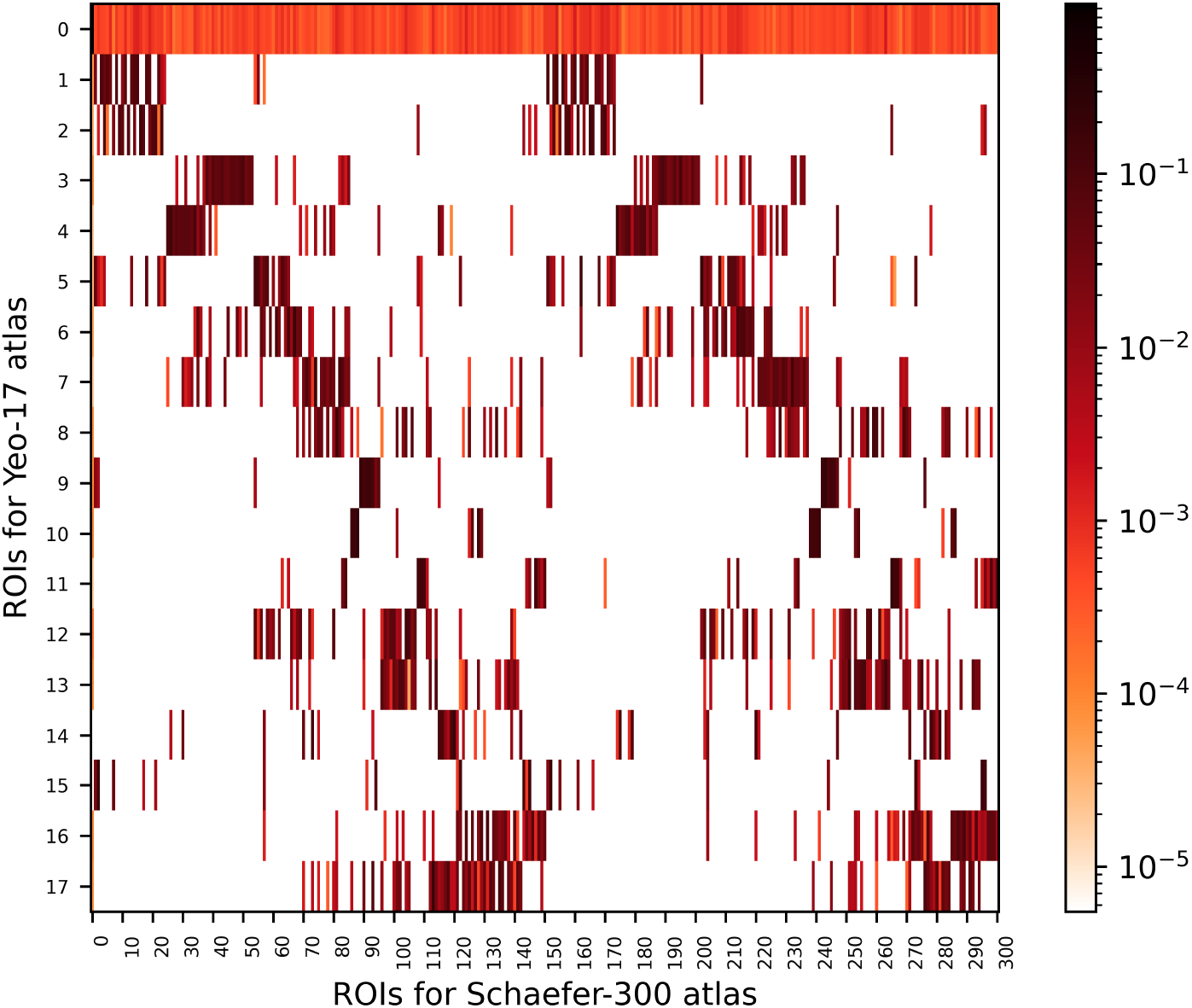
Dice Score Map between the Yeo-17 Networks atlas and the 300 parcellation Schaefer atlas. In a Dice Score Map, the larger the Dice score, the larger the percentage of overlap. Due to Schaefer’s larger quantity of ROIs, several different ROIs overlap a single Yeo-17 ROI. In this Dice Map, the 0 ROI for Yeo-17 represents the background of the image, or the empty space in the image. This ROI not having a Dice value of 0 indicates that both atlases don’t cover the same amount of brain volume.

### Adjusted Mutual Information

Using another python script, provided in Neuroparc GitHub repository, the Adjusted Mutual Information (AMI) between atlases was calculated. See the Adjusted Mutual Information section in Methods for more information about the calculation. The results, displayed in Figure 3, affirm the necessity for having access to multiple different parcellation methods during data analysis. Each atlas was created using different reference data and designed to track specific structure or functionality present in the brain. If atlases were similar to the point of being interchangeable, this wide variety of AMI scores would not exist, along with the reason for having a repository of atlases. The AMI amongst the Schaefer atlas set [32], DS atlas set [39], and Slab [40] atlases was consistently greater than 0.8, an expected result due to their creation using the same methodologies with different parameters and tolerances.

**Figure 3:**
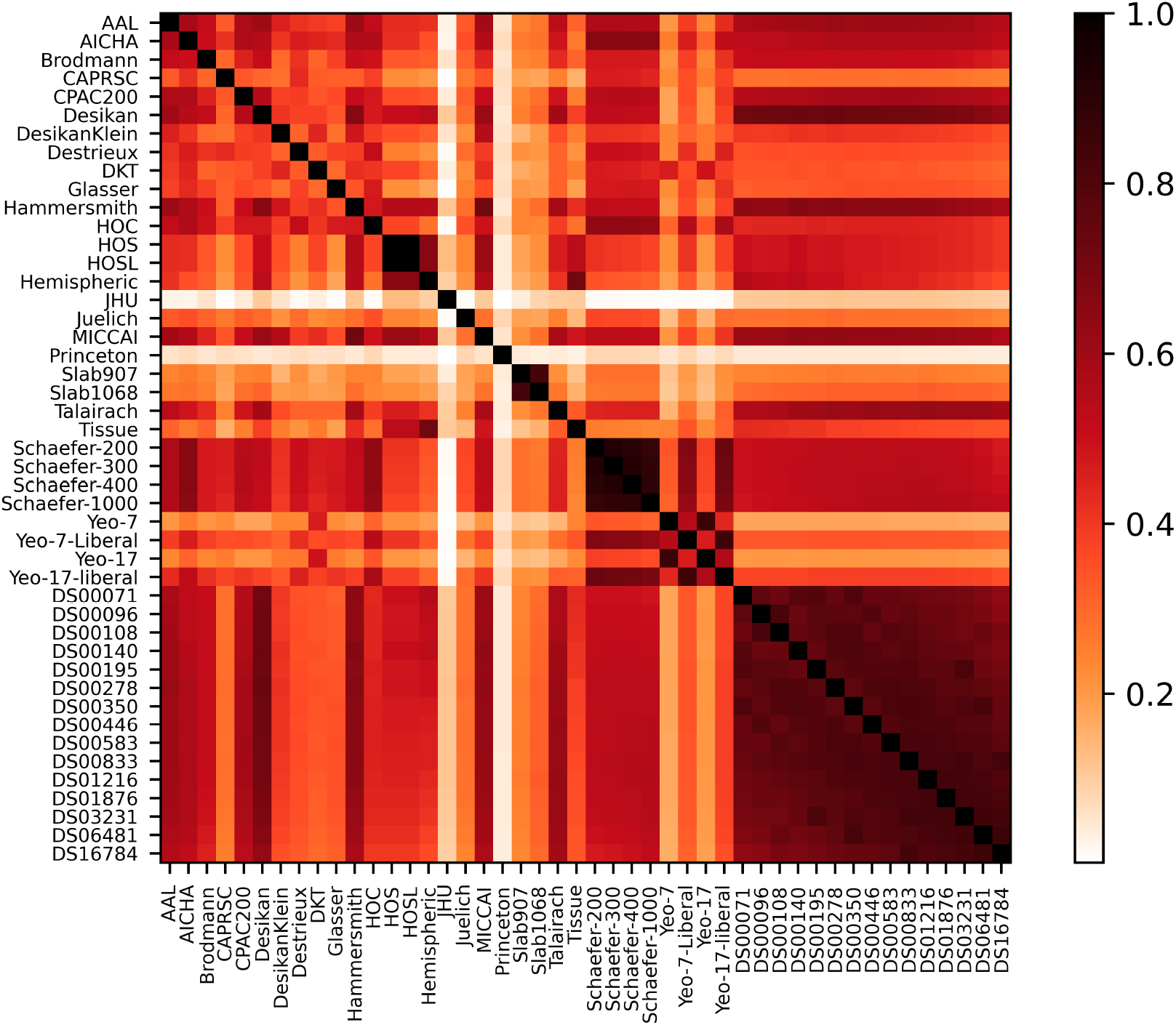
The adjusted mutual information between atlases contained within Neuroparc. Atlases that were generated from the same algorithm using different parameters, such as Yeo, Slab, Schaefer, and DS have an expected high amount of mutual information.

Both the JHU [34] and Princeton [35] atlases displayed a consistently low (< 0.3) AMI value for the majority of other atlases. This can be explained by the fact that both atlases only relate to anatomical sub-structures, such as the visual cortex for JHU and hippocampal region for Princeton. Their limited coverage of the brain results in less mutual information with the other surface-based or volume-based atlases.

## 3 Discussion

### Why use Neuroparc?

The purpose of the Neuroparc atlas collection and metadata formatting method is two fold: (1) to provide a repository of standardized parcellations that can be used interchangeably without any additional effort, and (2) to document all relevant information about each parcellation for easy use in research. Neuroparc succeeds in both of these aspects, as well as enabling a new level of comparison between atlases. Using the formatting method proposed in this paper, any user of the repository has the ability to find where each atlas came from, how many different ROIs exist in the atlas, the location and size of each of each ROI, how the segmentation of an atlas compares to others, and whether there is a significant correlation between areas covered by ROIs from different atlases. The formatting method also allows for constant improvement and refinement of atlas metadata, discussed below in Future Development. By standardizing the atlases, researchers can easily analyze MRI data in MNI space using a variety of atlases without additional processing. Metrics provided by Neuroparc, such as adjusted mutual information and the dice coefficient, also inform users as to how the atlases are related.

### Potential Issues

The method used to generate the atlases for Neuroparc can result in the loss of ROIs due to down-sampling and registration. The chance of this occurring inversely correlates with the average size of ROIs in a given atlas. The ROIs that are lost are still cataloged in the corresponding JSON file, with a value of “null” being given for the center coordinates and size. While there do exist ways to attempt to prevent this loss of information, excessive manipulation of a given atlas depending on the voxel size potentially compromises any conclusions derived using said atlas. As such, the parcellations in Neuroparc do not incorporate these additional methods and it us on the user to decide how best to resample the corresponding 1mm voxel parcellation to fit their unique needs.

### Future Development

With the current iteration of Neuroparc there are several routes for improvement. The most apparent is the expansion of the atlas collection. Our proposed methodology for standardizing new atlases and tracking metadata makes this task a simple one. With the emphasis on clear and concise information, approval of any new set of atlases is a quick and simple process. There also exists the ability to standardize all atlases to other spaces besides MNI, allowing for atlases offered in both different voxel sizes and standardized spaces. Another route for growth of Neuroparc is the anatomical labeling of atlases whose ROIs do not have clearly defined anatomical boundaries. The anatomical labels currently in Neuroparc are taken from the published work where they were first made. To keep the rational of the original authors, very little was done to the labels provided, mainly rewording for clarity and to follow the largest structure to smallest structure method. Due to subjective nature of labeling the atlases, an agreed upon anatomical labeling reference would first have to be made. From there, anatomical labeling could be assigned pragmatically.

The methodology proposed in this paper, as well as the atlas repository in Neuroparc, attempt to address the lack of standardization and centralization for brain atlases. We believe that Neuroparc exemplifies the first steps towards solving this issue. We call on other researchers to utilize the resources contained within and encourage everyone to contribute.

## 4 Methods

### Data Compilation

The atlases contained in Neuroparc were collected from a variety of locations. As previously noted, there is no current standard for atlas storage, so all gathered datasets are converted into a single format. Collected atlases were re-sampled using AFNI’s 3dresample [37] to either 1 mm^3^, 2 mm^3^, or 4 mm^3^ voxel resolution and then registered to a reference T1-weighted image described below. The sources and additional information for each of the atlases can be found in the README file in the GitHub repository.

### Reference Brain

To allow direct comparison between different atlases, a standard reference brain must be used for all involved atlases. Within Neuroparc, a single reference brain is provided and multiple resolutions, yielding a consistent coordinate space. Neuroparc uses Montreal Neurological Institute 152 Nonlinear 6th generation reference brain, abbreviated MNI152NLin6 in the file naming structure [41]. While there are a symmetric and asymmetric version of the MNI152NLin6 T1-weighted image, Neuroparc atlases are registered to the symmetric version, which is used by both FSL 5.0 [38] and new versions of SPM. However, code provided in Neuroparc allows for the registering of any atlas to any reference brain the user chooses.

This image is stored in a GNU-zipped NIfTI file format of a T1-weighted MRI and is available in Neuroparc at three resolutions (1 mm^3^, 2 mm^3^, and 4 mm^3^) for easy use when registering. The naming convention for these files clearly displays their source and resolution as: MNI152NLin6_res-<*resolution*>_T1w.nii.gz. For example, the format of the *resolution* input would be “1×1×1” for the 1 mm^3^ resolution.

### Atlas Images and Processing

The atlas images compiled in Neuroparc were stored in the form of GNU-zipped NifTi files containing the parcellated atlas. In these files, each region of interest (ROI) within the parcellated image is denoted by a unique integer ranging from 1 to n, where n is the total number of ROIs. Atlases were resampled to the desired voxel resolution through the use of AFNI’s 3dresample [37] and then registered to the MNI image of the same resolution using FSL’s flirt function [38]. The naming convention for the resulting atlas was: <*atlas_name*>_space-MNI152NLin6_res-<*resolution*>.nii.gz. The *atlas_name* field is unique for each atlas image, ideally no more than two words long without a space in between (e.g. Yeo-17 [36], Princetonvisual [35], HarvardOxford [31]).

### Atlas Metadata

Using a Python script in Neuroparc, a JSON file containing relevant meta-data was generated for each of the atlases. This file was split into two sections: region-wide and atlas-wide information. The naming convention of the JSON file follows that of the atlas image, with an atlas with 1 mm^3^ voxel size being named <*atlas_name*>_space-MNI152NLin6_res-1×1×1.json.

The term “region-wide” refers to information unique to each region of interest (ROI) in said atlas. This information includes the voxel value for that ROI in the atlas, an anatomical label (if possible), the coordinates of the center of the ROI, and the number of voxels that make up that region. The center and size can be calculated using provided code in the scripts directory of Neuroparc.

Although the label must be specified, this information is not relevant for all atlases. For example, atlases that are generated using algorithmic means, like Slab [40], have ROIs that do not strongly correlate to individual anatomical regions. In that case, NULL should be used for the labels of the regions. For ROIs that do have anatomical significance, the naming should follow a hierarchical format in order of largest region to smallest with an underscore in between each name. Modifiers, such as “Superior” or “Medial” can be placed before the anatomical region. An example of this is in the Desikan atlas [27], which contains the region with label “L _rostral _anterior _cingulate_cortex”. The main purpose of labeling is to clearly convey the location of an ROI and any anatomical significance it may have. The avoidance of unique abbreviations or terminology that is not widely used helps with the ease of use for individuals new to MRI analysis.

Figure 4 shows an example json file.

**Figure 4:**
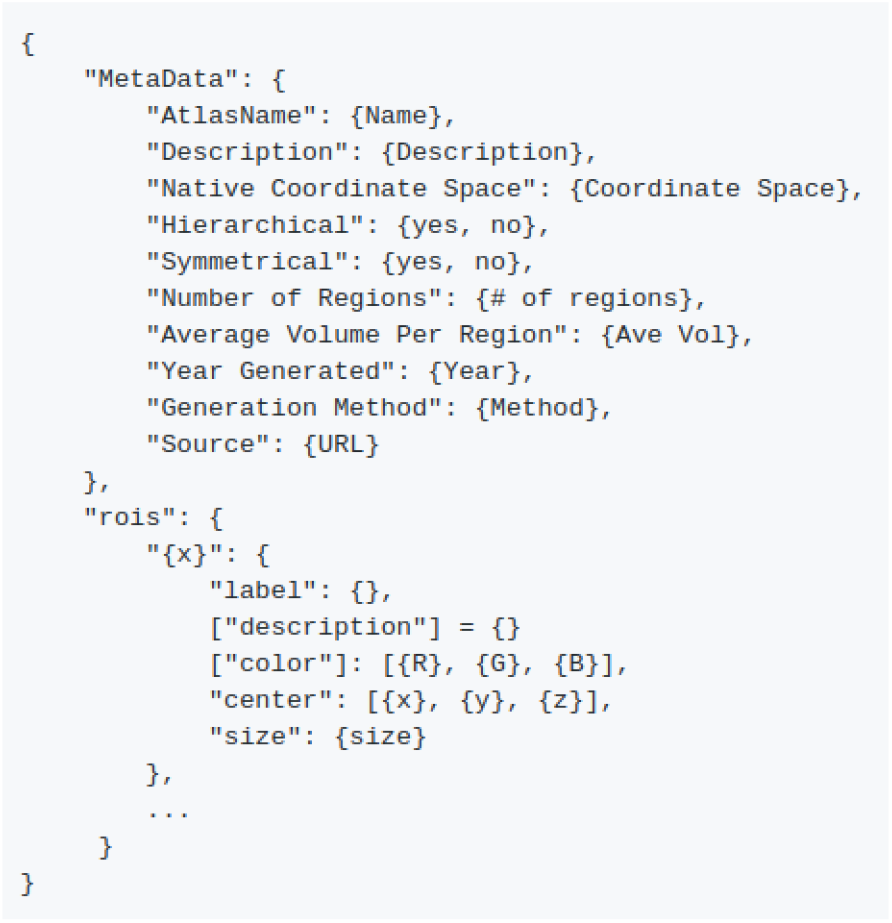
An example JSON file rubric for storing atlas metadata. Metadata stored in brackets (“color”, “description”, etc.) is optional but encouraged.

Optional fields in the region-wide data include description and color. Description can be used to provide more information than the region label if necessary. An example of this use is in the Yeo-7 Networks atlas [36]. The label for this atlas is in the form ‘7Networks_2’, but the description for that label is the corresponding functional network, ‘Somatomotor’ in this case. The color field must be given in the form [R, G, B] and is only used if the user wants to specify the colors of the regions upon visualization.

Brain-wide data must include the name, description, native coordinate space, and source of the atlas. The name field allows for more elaboration than in the name of the file. The description is more flexible, allowing the creator of an atlas to briefly describe important information for users of their atlas. The intended use case or the method of generation are examples of information provided in this field. Since all atlases in Neuroparc are stored in the same coordinate space, the coordinate space used during the creation of the atlas must be specified.

Finally, the publication detailing the atlas should be included in the source field so users can have a more full understanding of the atlas being used. Optional fields for brain-wide data can all be calculated, including the number of regions, the average volume per region, whether the segmented regions are hierarchical, and if the atlas is symmetrical.

The full description and format of the atlas specification is available within Neuroparc at https://github.com/neurodata/neuroparc/tree/master/atlases/atlas_spec.md.

### Dice Coefficient

As a way to compare the different atlases to each other, since each has been registered to MNI space, we calculated the Dice Coefficient between atlases. The Dice coefficient is a measure of similarity between two sets [42]. Specifically, it measures a coincidence index (CI) between two sets, normalized by the size of the sets. Let *h* be the number of points overlapping in the sets *A* and *B*, and *a* and *b* are the sizes of their corresponding sets. If the two sets are labelled regions in segmented images, then the Dice coefficient between any pair of regions between the images is given by

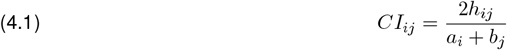

where *i* is the region in image 1 and *j* is the region in image 2. The result is a similarity matrix, as shown in Figure 2. Since this map visualizes similarity between two regions in two atlases, the information provided by the Dice map can be used quantify which regions in a given atlas are most similar to regions in another atlas. This method has proven valuable for performing inference with parcellations lacking anatomical annotation, as it allows conclusions realized at the parcel level to be inferred at the anatomical level [43].

### Adjusted Mutual Information

Adjusted mutual information is another measure of the similarity of two labelled sets, quantifying how well a particular point can be identified as belonging to a region given another region. It differs from the Dice coefficient in that it tends to be more sensitive to region size and position relative to other measures. [44]

Similar to the Dice coefficient, Adjusted mutual information is not dependent on a region’s label [45]. Volumes that share many points are likely to be have a higher mutual information score all else being equal [46].

To assure that all atlas comparisons were on the same scale, Neuroparc computes the adjusted mutual information score. Let *H*(·) denote entropy, *N* be the number of elements (voxels) in total, and *E*(*MI_A_B*) denote the expected mutual information for sets of size a and b. Here, *P_A_*(*i*) is the probability that a point chosen randomly from the set A will belong to region i. [47]

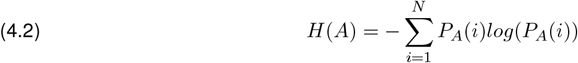

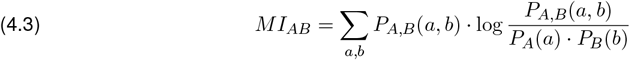

where *P_A,B_*(*a, b*) is the probability that a voxel will belong to region a in set A and region b in set B.

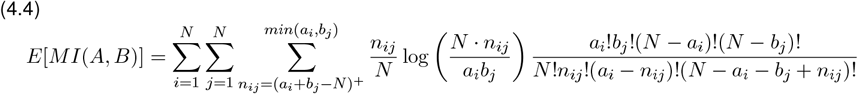

where *a_i_* is the number of voxels in region i of set A and *b_j_* is the number of voxels in region j of set B. (*a_i_* + *b_j_* − *N*)^+^ = *max*(1, *a_i_* + *b_j_* − *N*).

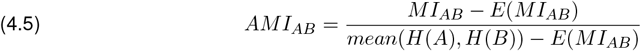

as provided in [48].

Figure 3 shows the adjusted mutual information between all pairs of atlases. The information provided for this score is atlas-wide, while the Dice score was computed per region to generate a map. The similarity between groups of atlases, such as the various Schaefer atlases, the Yeo liberal atlases, and the DS atlases, is immediately apparent. Recent work highlighting the complementary information provided by disparate parcellations stresses the importance of the availability and ease-of-use of a collection of parcellations from heterogeneous sources [43, 49–51].

## 5 Data Availability

All atlases and scripts described in this paper are available through a Github repository https://github.com/neurodata/neuroparc. A more extensive repository can be found in an OSF repository https://osf.io/67a3t/, which also contains all Dice and AMI figures and adjacency matrix results. A link is also available from https://neurodata.io/mri.

## 6 Code Availability

Code for processing is publicly available and can be found on GitHub under the scripts folder (https://github.com/neurodata/neuroparc). Examples of useful functions include resampling parcellations to a desired voxel size, the ability to register parcellations to any given reference image, and center calculation for regions of interest for 3D parcellations. Jupyter notebook tutorials are also available for learning how to prepare atlases for being added to Neuroparc. All code is provided under the Apache 2.0 License.

Visualizations are generated using both MIPAV 8.0.2 and FSLeyes 5.0.10 to view the brain volumes in 2D and 3D spaces [52, 53]. Figure 1 can be created using MIPAV triplanar views of each atlases with a striped LUT.

## Acknowledgements

We would like to acknowledge generous support from National Science Foundation (NSF) under NSF Award Number EEC-1707298. Reasearch was partially supported by funding from Microsoft Research.

## Author contributions

R.L., E.W.B., P.M., G.A., D.A.P., and J.T.V. prepared the data source and performed the comparative data analysis. S.R., P.F, A.L., and A.N. performed the technical validation. All authors prepared the manuscript.

